# PROTEIN CODING VARIATION IN OUTBRED LABORATORY MOUSE STOCKS PROVIDES A MOLECULAR BASIS FOR DISTINCT RESEARCH APPLICATIONS

**DOI:** 10.1101/2022.07.26.501579

**Authors:** Belinda Cornes, Carolyn Paisie, Emily Swanzey, Andrew Schile, Kelly Brackett, Laura Reinholdt, Anuj Srivastava

## Abstract

Outbred laboratory mice (*Mus musculus*) are readily available and have high fecundity, making them a popular choice for biomedical research, especially toxicological and pharmacological applications. Direct high throughput genome sequencing (HTS) of these widely used research animals is an important genetic quality control measure that ensures research reproducibility. HTS data have been used to confirm the common origin of outbred stocks and to molecularly define distinct outbred populations. But these data have also revealed unexpected population structure and homozygosity in some populations; genetic features that emerge when outbred stocks are not properly maintained. We used exome sequencing to discover and interrogate protein coding variation in a newly established population of Swiss-derived outbred stock (J:ARC) that is closely related to other, commonly used CD-1 outbred populations. We used these data to describe the genetic architecture of the J:ARC population including heterozygosity, minor allele frequency, LD decay, and we defined the novel, protein-coding sequence variation. These data reveal the expected genetic architecture for a properly maintained outbred stock and provide a basis for on-going genetic quality control. We also compared these data to protein coding variation found in a multiparent outbred stock, the Diversity Outbred (J:DO). We found that the more recently derived, multiparent outbred stock has significantly higher interindividual variability, greater overall genetic variation, higher heterozygosity, and fewer novel variants than the Swiss derived J:ARC stock. However, among the novel variants found in the J:DO stock, significantly more are predicted to be protein-damaging. That individuals from this population can tolerate a higher load of potentially damaging variants highlights the buffering effects of allelic diversity and the differing selective pressures in these stocks. While both outbred stocks offer significant individual heterozygosity, our data provide a molecular basis for their intended applications, where the J:DO are best suited for studies requiring maximum, population-level genetic diversity and power for mapping, while the J:ARC are best suited as a general-purpose outbred stock with robust fecundity, relatively low allelic diversity, and less potential for extreme phenotypic variability.

## INTRODUCTION

There are 1,374 inbred strains and outbred laboratory mouse stocks cited in the biomedical research literature (Bult *et al*. 2019). Genome sequencing of inbred strains has confirmed the expected homozygosity and intrastrain genetic divergence that are the results of decades of inbreeding and reproductive isolation (Mouse Genome Sequencing *et al*. 2002; Keane *et al*. 2011; Yalcin *et al*. 2011; Chang *et al*. 2017; Srivastava *et al*. 2017; Lilue *et al*. 2018). Because inbred mice are genetically identical, interindividual phenotypic variability can be attributed to experimental variables and to a lesser extent, epigenetic variation (Weichman and Chaillet 1997). These unique features make inbred mouse strains a popular choice for biomedical researchers who endeavor to minimize genetic variability and to quantify experimental (extrinsic, non-genetic) variability. However, inbred strains do not recapitulate the extent of interindividual variation found in a human cohort, patient group, or population making them inappropriate for studies that seek to model such variation.

In contrast to inbred strains, outbred stocks exhibit interindividual variation and heterozygosity, coupled with higher fecundity. The four most widely cited outbred stocks are CF-1, Swiss Webster, NMRI, and CD-1 (Bult *et al*. 2019), but all trace their origins to a colony at The Rockefeller Institute that was established with 2 male and 7 female ‘Swiss’ mice imported from the Pasteur Institute in 1926 (Chia *et al*. 2005). Over the ensuing decades, mice from these stocks were imported by commercial breeders, NIH, and academic institutions where colonies were ideally maintained with at least 25 breeding pairs to maintain a coefficient of inbreeding (F) of less than 1% per generation with selection for maximum fecundity (Festing *et al*. 1972) (Chia *et al*. 2005). A variety of approaches have been used to examine the genetic variation resident in these outbred stocks including alloenzyme analysis and other biochemical approaches (Groen and Lagerwerf 1979; Rice and O’Brien 1980; Cui *et al*. 1993), high-density SNP panels (Aldinger *et al*. 2009; Yalcin *et al*. 2010), and most recently low coverage genome sequencing (Yalcin *et al*. 2010; Nicod *et al*. 2016). Data from these studies provide estimates of minor allele frequency, heterozygosity, and coefficients of inbreeding. Specifically, low coverage sequencing of these stocks has revealed 1) relatively low allelic diversity given their common origin, 2) appreciable levels of heterozygosity of common variants (0.20-0.35) in properly maintained populations, and 3) relatively few novel variants (<5%) relative to the C57BL/6J reference genome and other common inbred laboratory strains (Yalcin *et al*. 2010; Nicod *et al*. 2016).

The JAX Swiss Outbred stock was imported to The Jackson Laboratory (JAX) from The Animal Resources Centre (ARC) in Canning Vale, Western Australia in 2020 as a group of 128 individuals (64 male and 64 female). To fully characterize the extent of protein coding variation in this new outbred population, we sequenced the exomes of the 32 male and 32 female G3 animals that were used to initiate new breeding funnels at JAX (J:ARC). We used these data to estimate inbreeding, heterozygosity, and interindividual variation, and to molecularly define the colony. These metrics confirmed the outbred nature of this new colony of CD-1 stock, and additionally reveal 5,105 (3.8%) novel protein coding and splice site variants, which falls in the range of the proportion of novel variants found in other outbred colonies and stocks (Yalcin *et al*. 2010; Nicod *et al*. 2016). To provide a molecular genetic quality control standard for outbred stocks, we compared the J:ARC exome call sets to exome variant call sets from very different outbred stock, derived from a multiparent outbred population, the Diversity Outbred (J:DO). We found that the J:DO have higher population level genetic variation (~3.6X), higher interindividual variation, higher heterozygosity, and higher allelic diversity due to their multi-substrain parental origins. We found fewer novel variants in the J:DO, yet these novel variants are predicted to have higher impact on protein function. These common and distinct genomic features of the J:ARC and J:DO can serve as the basis for estimating sample size, where fewer J:ARC mice are needed to capture the full range of segregating variation in the population, but a similarly sized cohort of J:DO mice offers more interindividual genetic variation and higher population level genetic variation. Both types of outbred stocks can be used for genetic mapping, however, their distinct genetic architectures must be accounted for in study design as previously shown (Gatti *et al*. 2014; Nicod *et al*. 2016), and should be regularly monitored to ensure reproducibility.

## MATERIALS & METHODS

### Samples

#### J:ARC(S), JAX Swiss Outbred

The JAX Swiss Outbred stock was imported to The Jackson Laboratory from The Animal Resources Centre (ARC), Canning Vale, Western Australia in 2020. The imported mice (G0) were paired into 64 breeding units, and sperm and eggs were harvested from the G1 offspring. To establish each of the 32 breeding funnels for live colony maintenance in the JAX vivarium barrier, two distinct units were selected for reciprocal *in vitro* fertilization and IVF-generated embryos were pooled. The resulting live-born animals from each funnel were designated G2 (Figure 1). These mice were subsequently bred according to the Poiley rotational breeding scheme to produce 32 breeder pair units / funnels at G3(Poiley 1960). Spleen samples were collected the 64 G3 J:ARC (J:ARC(S), RRID:IMSR_JAX:034608, JAX stock number 034608) (32 females and 32 males) that populated each unit (Supplementary Data 1).

**Figure 1.**
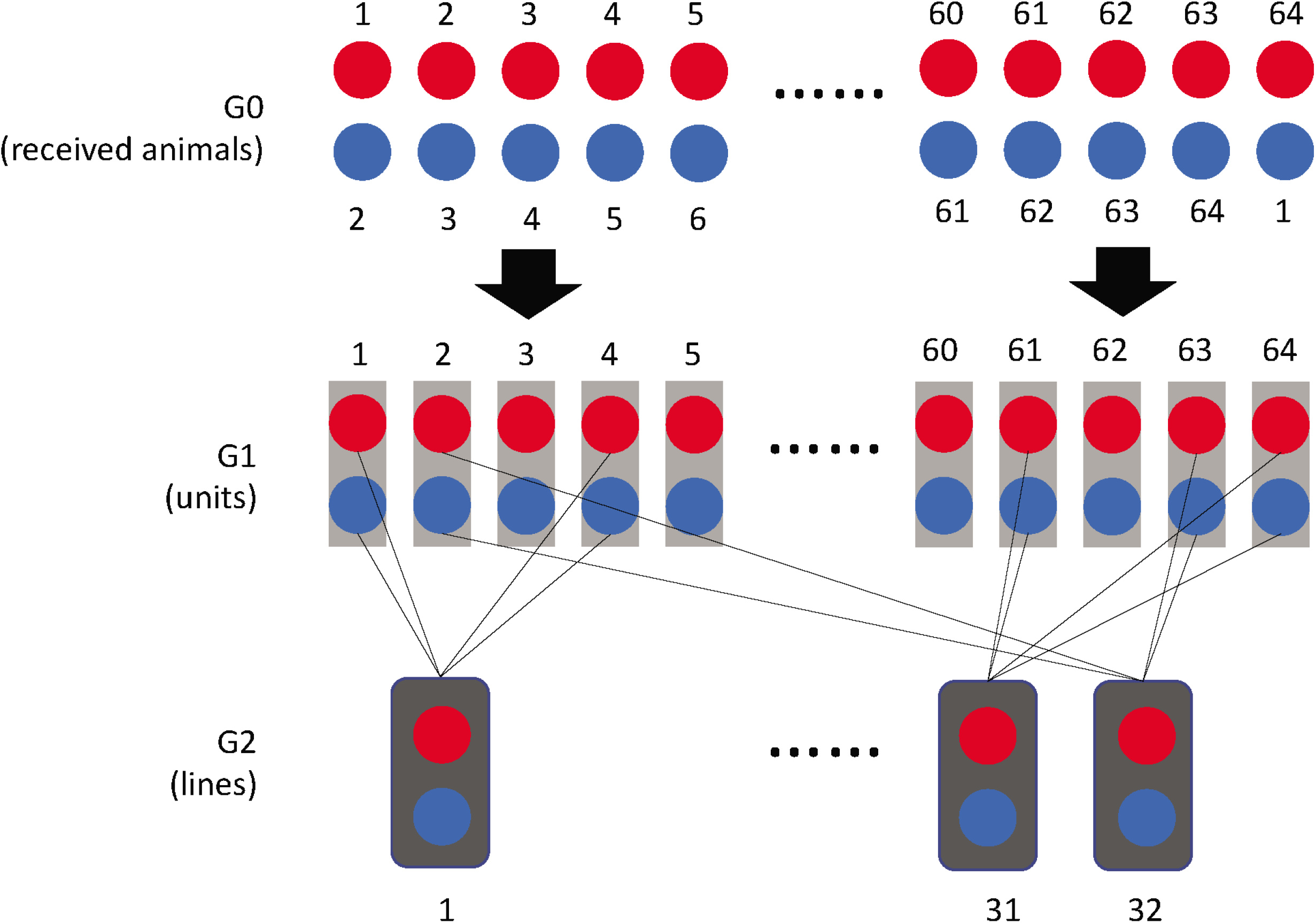
The Poiley method used for the importation and breeding of the JAX Swiss Outbred (J:ARC) population. G0 were live animals from The Animal Resources Centre (ARC) in Canning Vale that were subsequently bred and rederived through IVF to create 32 distinct breeding lines at The Jackson Laboratory which continue to be maintained according to the Poiley method to minimize inbreeding.

#### J:DO, JAXDiversity Outbred

We collected spleen from 20 J:DO, Diversity Outbred (RRID:IMSR_JAX:009376, JAX stock #9376) mice from The Jackson Laboratory, 10 female and 10 male, non-sibling mice, breeding generation 43 (Supplementary Data 1).

### DNA Isolation, Exome Library Preparation, and Sequencing

DNA isolation, exome library preparation, and sequencing were performed by Genome Technologies at The Jackson Laboratory. DNA was isolated from spleen using the NucleoMag Tissue Kit (Machery-Nagel) according to the manufacturer’s protocol. DNA concentration and quality were assessed using the Nanodrop 8000 spectrophotometer (Thermo Scientific), the Qubit Flex dsDNA BR Assay (Thermo Scientific), and the Genomic DNA ScreenTape Analysis Assay (Agilent Technologies). Mouse exome libraries were constructed using the KAPA HyperPrep Kit (Roche Sequencing and Life Science) and SureSelect XT Community Design Mouse All Exon v2 Target Enrichment System (Agilent Technologies), according to the manufactures’ protocols. Briefly, the protocol entails shearing the DNA using the E220 Focused-ultrasonicator (Covaris), size selection targeting 400 bp, ligating Illumina specific adapters, and PCR amplification. The amplified DNA libraries are then hybridized to the Mouse All Exon probes (Agilent Technologies) and amplified using indexed primers. The quality and concentration of the libraries were assessed using the High Sensitivity D1000 ScreenTape (Agilent Technologies) and KAPA Library Quantification Kit (Roche Sequencing and Life Science), respectively, according to the manufacturers’ instructions. Libraries were sequenced 150 bp paired-end on an Illumina NovaSeq 6000 using the S4 Reagent Kit v1.5.

### Sequence Data Analysis

An overview of the sequence data analysis pipeline is shown in Supplementary Figure 1. All reads were subjected to quality control using an in-house QC script. Samples with base qualities greater ≥ 30 over 70 % of read length were used in the downstream analysis. High-quality reads were mapped to the mouse genome (build-mm10) using BWA-mem (bwa-0.7.9a) at default parameters (Aldinger *et al*. 2009). The resulting alignment was sorted by coordinates and converted to binary alignment map (BAM) format by Picard v 2.8.1-SortSam utility (http://picard.sourceforge.net). Picard-MarkDuplicates module was used to remove the duplicates from data. The Genome Analysis Toolkit (GATK v4) (McKenna *et al* 2010; DePristo *et al* 2011) module BaseRecalibrator was used to pre-process the alignments. Target capture efficiency was determined using Picard-HsMetrics (1.95). The recalibrated bam alignment file was used to input GATK-Haplotype Caller at parameters -stand_call_conf 50.0, - stand_emit_conf 30.0 and variants calls were restricted to the target region (Mouse All Exon v2). Finally, raw variants calls were soft filtered using GATK VariantFiltration (DP < 25, QD < 1.5 and FS > 60), annotated by snpEff 3.6.c (Cingolani *et al* 2012) and the highest impact variant was reported by GATK VariantAnnotator. All variants were further annotated with mouse dbSNP v150.

GATK-HaplotypeCaller in GVCF mode was used for joint genotyping of J;DO, J:ARC, and combined (J:DO and J:ARC) samples. The CombineGVCF utility was used to gather all the samples and then executed the GenotypeGVCF command. All analyses were performed on jointly called variants and only included variants found in both “J:ARC only” and “J:DO only” called datasets (N: 120,864). This effectively revealed the differences between the two datasets and directly provided the data needed to resolve genotypes.

### Heterozygosity

To observe the distribution of mean heterozygosity across all samples, Plink was used (Purcell *et al*. 2007). The ‘--het’ option in Plink was used to create heterozygous information for each sample which included the observed number of homozygous genotypes ‘[O(Hom)]’ and the number of non-missing genotypes ‘[N(NM)]’. This information was then used to calculate the observed heterozygosity rate per individual using the formula ‘(N(NM) - O(Hom))/N(NM)’.

To calculate per variant heterozygosity for variants shared between J:DO and J:ARC, the number of samples in which a shared variant was heterozygous was divided by the total number of genotyped samples. The inbreeding coefficient (F_H_), was calculated using *VCFtools* (−het) as previously described (Stoffel *et al*. 2016; Foster *et al*. 2021). PLINK was used to calculate *r^2^* for all pairs of autosomal SNPs called from joint genotyping in the J:ARC samples (425,409) and the J:DO samples (117,429). SNPs that were missing in more than 5% of the samples and that were monomorphic were removed. The 95^th^ percentile of *r^2^* values for SNPs spaced up to 1 Mb apart were selected.

### Allele Frequency

To observe differences between allele frequencies across the genome between J:DO and J:ARC samples, the minor allele frequency (MAF) was calculated separately for J:DO and J:ARC samples at each variant site using the ‘--freq’ option in Plink.

### Clustering/Dendrogram

Principal Component Analysis (PCA) plots were performed in Plink. PCA is a multivariate statistical method used to produce any number of uncorrelated variables (or principal components) from a data matrix containing observations across a number of potentially correlated variables. The principal components are calculated so that the first principal component accounts for as much variation in the data as possible in a single axis of variation (component), followed by additional components. Here the variants called in both J:ARC and J:DO datasets were used to observe the similarities within and between datasets.

To show similarity groups among the J:ARC samples, a dendrogram was performed using the ggdendro package in R-3.6.2 (https://cran.r-project.org). A dendrogram is a tree diagram showing hierarchical clustering that represents the relationships of similarity among a group of entities.

### Novel Variants

The joint genotyping variant call file (vcf) of J:ARC and J:DO were flagged for known variants in dbSNP150 (Sherry *et al*. 2001), European Variation Archive (EVA) (Cezard *et al*. 2022), Sanger mouses genome project (Keane *et al*. 2011) (ftp://ftp-mouse.sanger.ac.uk//REL-2004-v7-SNPs_Indels/mgp_REL2005_snps_indels.vcf.gz), mouse Collaborative Cross genome (ftp://ftp.sra.ebi.ac.uk/vol1/ERZ460/ERZ460702/Merged_69_flagged.tab.vcf.gz) (Srivastava *et al*. 2017), and Diversity Outbred low pass sequencing (in-house resource) using vcftools ‘vcf-annotate’ (Keane *et al*. 2011) and snpsift_4.2 (Cingolani *et al*. 2012). We also flagged J:ARC variants for their presence in J:DO joint and single sample variant call sets and did a similar analysis for J:DO variants. The variants not present in any of these resources were considered novel and were further annotated by snpeff v4.3 (Cingolani *et al*. 2012) using the mouse GRCm38.75 snpeff database. Finally, highest effect variants are selected by gatk-3.6 VariantAnnotator (McKenna *et al*. 2010). Functional gene and pathway annotations (including KEGG, GO, UP, BIOCARTA, REACTOME) for the genes harboring novel high impact variants were compiled using DAVID Bioinformatics tools. DAVID Bioinformatics tools were then used to identify clusters of genes with related functional annotations using a medium classification stringency and a Benjamini score of >0.05 as the significance threshold for enrichment (Sherman *et al*. 2022)

## Results and Discussion

The J:ARC outbred stock currently maintained at The Jackson Laboratory was established through the importation of mice from the Australian Animal Resource Center (ARC) in 2020. The origins of this stock are CD-1 mice that were acquired by ARC from Charles River Laboratories in 1991 and 2005. Like most outbred laboratory mice, CD-1 mice trace their origins to 2 male and 7 female “Swiss” mice that were obtained by The Rockefeller Institute in 1926 (Chia *et al*. 2005). The live J:ARC colony was initiated with founder animals (G3) in 32 breeding funnels. We sought to determine the level of inbreeding and relatedness in the founder animals of this colony and to catalog the coding variants segregating in each funnel for future use in genetic quality control monitoring (Strobel *et al*. 2015). Since some outbred stocks differ quite dramatically, we sought to compare the genetic architecture of the outbred J:ARC to a multiparent outbred population (J:DO) to demonstrate the differing genetic architectures of these stocks and to establish a reference set of genetic variants for genetic monitoring. The J:DO mice are derived from eight founder inbred strains representing the three major subspecies of laboratory mice, *M.m.musculus, M.m.domestics,* and *M.m.castaneous.* This population captures 90% of the genetic variation in laboratory mice and with each generation of breeding, sufficient accumulated recombination to ensure high LD decay, and thereby maximum power for genetic mapping (Gatti *et al*. 2014; Saul *et al*. 2019). This population is maintained at the Jackson Laboratory through 175 breeding funnels using MateSel(Kinghorn and Kinghorn 2021) to minimize inadvertent phenotypic selection, to optimize diversity, and to prevent intracolony structure in each generation based on pedigreed population history.

### Genetic variation

We generated exome data from 64 J:ARC mice (32 male and 32 female) representing 32 breeding funnels at G3 and from 20 J:DO mice (10 male and 10 females) at generation 43. We generated ~203M high-quality reads on average, and after alignment to the reference genome, the mean target coverage was 162X (91% target covered at 30X) for J:ARC samples and 155X (90% target covered at 30X) for J:DO samples. Overall, 90% of the target exome regions were covered by 30 or more reads in both sets of samples (see Supplementary Figure 2).

Through joint genotyping, we identified 478,011 and 135,825 variants in J:DO and J:ARC samples, respectively, where a variant is defined as a SNP/INDEL present in a sequenced sample when compared to the C57BL/6J, inbred mouse reference genome (mm10, GRCm38). To adjust for the difference in sample size between the J:DO and J:ARC datasets and any potential influence this could have on overall discovery rates, we randomly subset the J:ARC samples to a set of twenty from joint genotyping results and then recorded the total number of variants present in the subset. We repeated this analysis over 100 iterations and found from this bootstrapping method that the J:DO harbor, on average, 3.6X more variants overall than J:ARC. This higher allelic diversity is expected in the J:DO since the stock is a multiparent population derived from eight diverse founder inbred strains.

### Heterozygosity and inbreeding

To evaluate heterozygosity in these two outbred populations, we estimated sample level heterozygosity for the entire set of variant calls (Figure 2A). The average, individual heterozygosity in J:ARC is 24%, which is consistent with previously estimates from low pass WGS for CFW outbred populations (Nicod *et al*. 2016) (Yalcin *et al*. 2010). In contrast, sample level heterozygosity in the J:DO samples was notably higher (nearly 40%) than J:ARC. Given that previous analyses have estimated J:DO haplotype-level heterozygosity at around 80% (https://www.jax.org/-/media/jaxweb/files/jax-mice-and-services/009376_J_DO_Mar2021.pdf), it is important to note that we used reference-based genotype calls and we calculated heterozygosity across the entire call set. Previously, heterozygosity for J:DO mice was calculated on a limited set of variants based on genotype probabilities of eight parental genotypes called from known polymorphic SNPs from the Mouse Universal Genotyping Array (MUGA) series (Morgan *et al*. 2015; Sigmon *et al*. 2020). To recapitulate this haplotype-based approach using our exome variant calls, we used 3007 calls at markers from GigaMUGA genotyping array (Morgan *et al*. 2015) and for this subset of variants, we found heterozygosity of ~82% consistent with previous array-based estimates (see Supplementary Figure 3).

**Figure 2.**
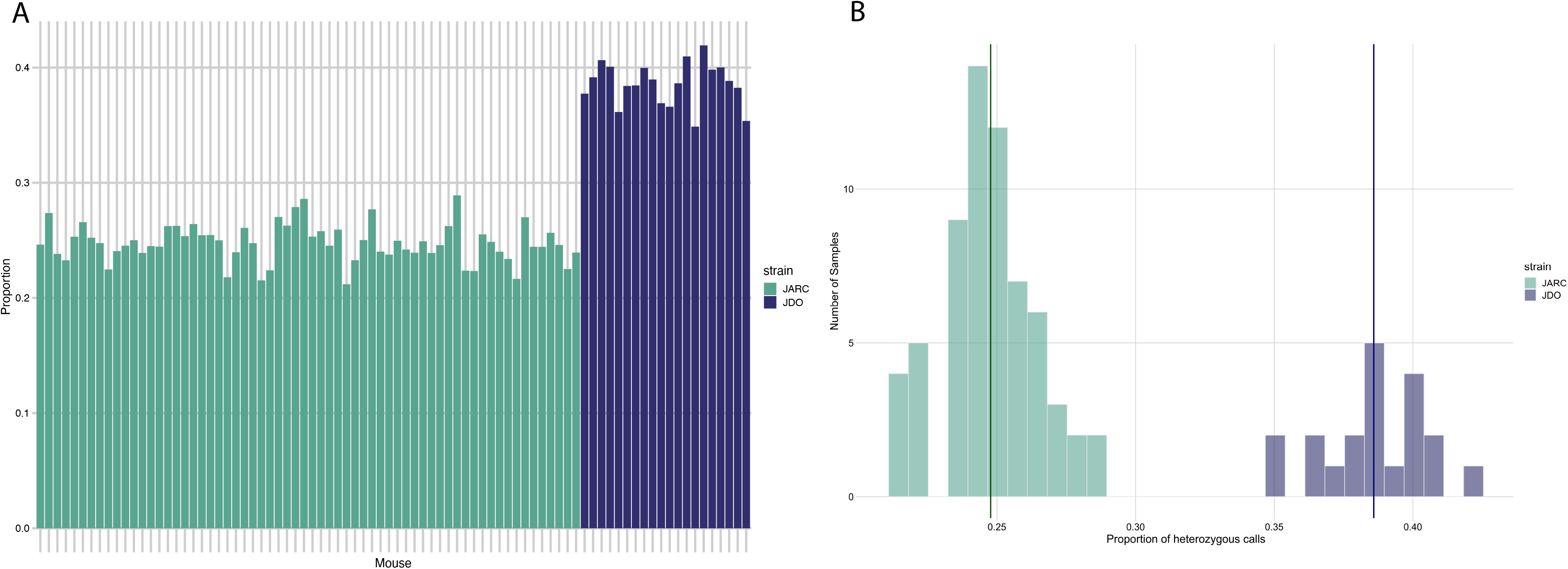
Heterozygosity in J:ARC (green) and J:DO (blue) datasets. The proportion (**A**) of heterozygous calls per samples and distribution of heterozygous calls (**B**) for J:ARC (green) and J:DO (blue) datasets. The mean heterozygosity for J:ARC samples was 0.25 while in J:DO it was 0.39. Samples from both datasets were joint called and this analysis included only those variants that are were called in both datasets (120,864)

To compare the variant level heterozygosity between the two populations, we identified the variants that are common between J:ARC and J:DO and then for each variant determined the number of samples with heterozygous genotypes (Figure 2B). We found that a lower proportion of J:ARC samples are heterozygous for any given variant than the J:DO, and that there is greater intraindividual variability in the proportion of heterozygous variants in the J:DO population. To further explore these differences in the distribution of heterozygosity, we calculated the minor allele frequencies (MAF) for each of the common variants across the genome (Figure 3A-C). We found that for each common variant, the MAF in the J:ARC was consistently lower (mean 0.18, median 0.13) than the J:DO (mean 0.28, median 0.28) and similar to previously published MAF for variants segregating in properly maintained outbred CD-1 mouse populations (Yalcin *et al*. 2010).

**Figure 3.**
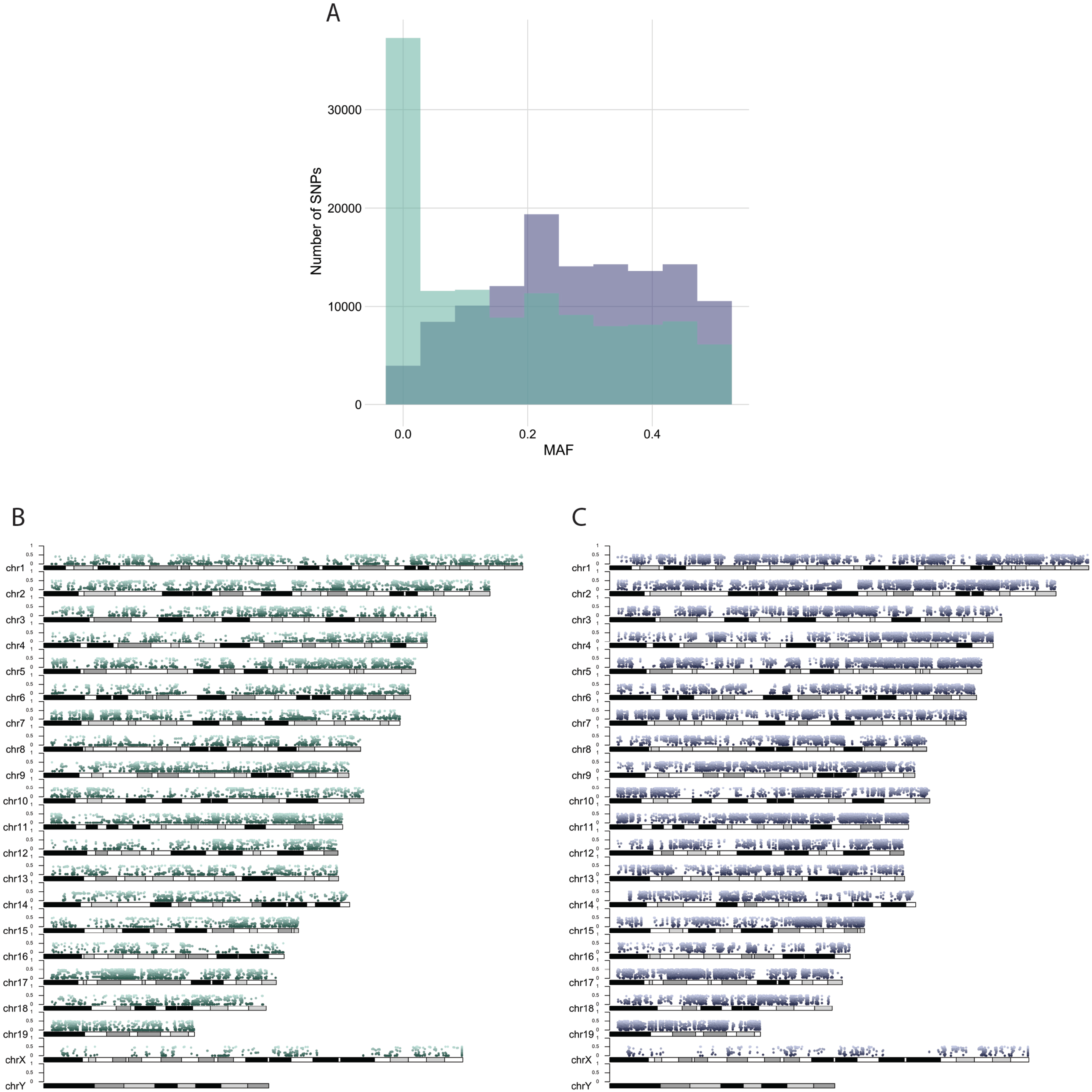
Allele Frequency in J:ARC (green) and J:DO (blue) datasets. Samples from both datasets were joint called and this analysis was restricted to those variants found in keeping both datasets (120,864). Allele frequency was calculated using Plink. (**A**) MAF distribution for 120,864 SNPs for J:ARC (green) and J:DO (blue). Monomorphic variants are not removed. (**B**) Allele frequency at the variant level for J:ARC samples (mean 0.18, median 0.13). (**C**) Allele Frequency for J:DO samples at the variant level (mean 0.28, median 0.28).

Variant data can also be used to estimate the coefficient of inbreeding (F), which is a metric that describes the distribution of variants in a population. According to the Hardy-Weinberg (HW) principle, F=0 for a given variant indicates that it is in HW equilibrium. By this measure, a fully inbred strain has an inbreeding coefficient of F=1.0 (100%). When calculated across all J:ARC variants and samples, the average level of inbreeding is −.033 (−3.3%), which falls within expectations for an outbred population (Yalcin *et al*. 2010). Though fewer samples were used to generate the J:DO dataset, the inbreeding coefficient calculated from the J:DO was closer to HW equilibrium at −0.007 (−0.7%). These F coefficients indicate that both populations are maximally outbred. The negative values are the result of heterozygous calls that exceed HW expectations which in these datasets could be due to erroneous genotype calls or low sampling of the populations. While the coefficients of inbreeding for these populations are low, there are regions of the genome where shared variants are fixed. These regions of homozygosity are distributed through the J:ARC and J:DO genomes (Figure 4). It is likely that some homozygous variant calls (regions of homozygosity, ROH) are the result of heterozygous deletions. Because our methodology doesn’t distinguish between potential hemizygous and true homozygous variant calls, we may be overestimating the extent of ROH in these genomes.

**Figure 4.**
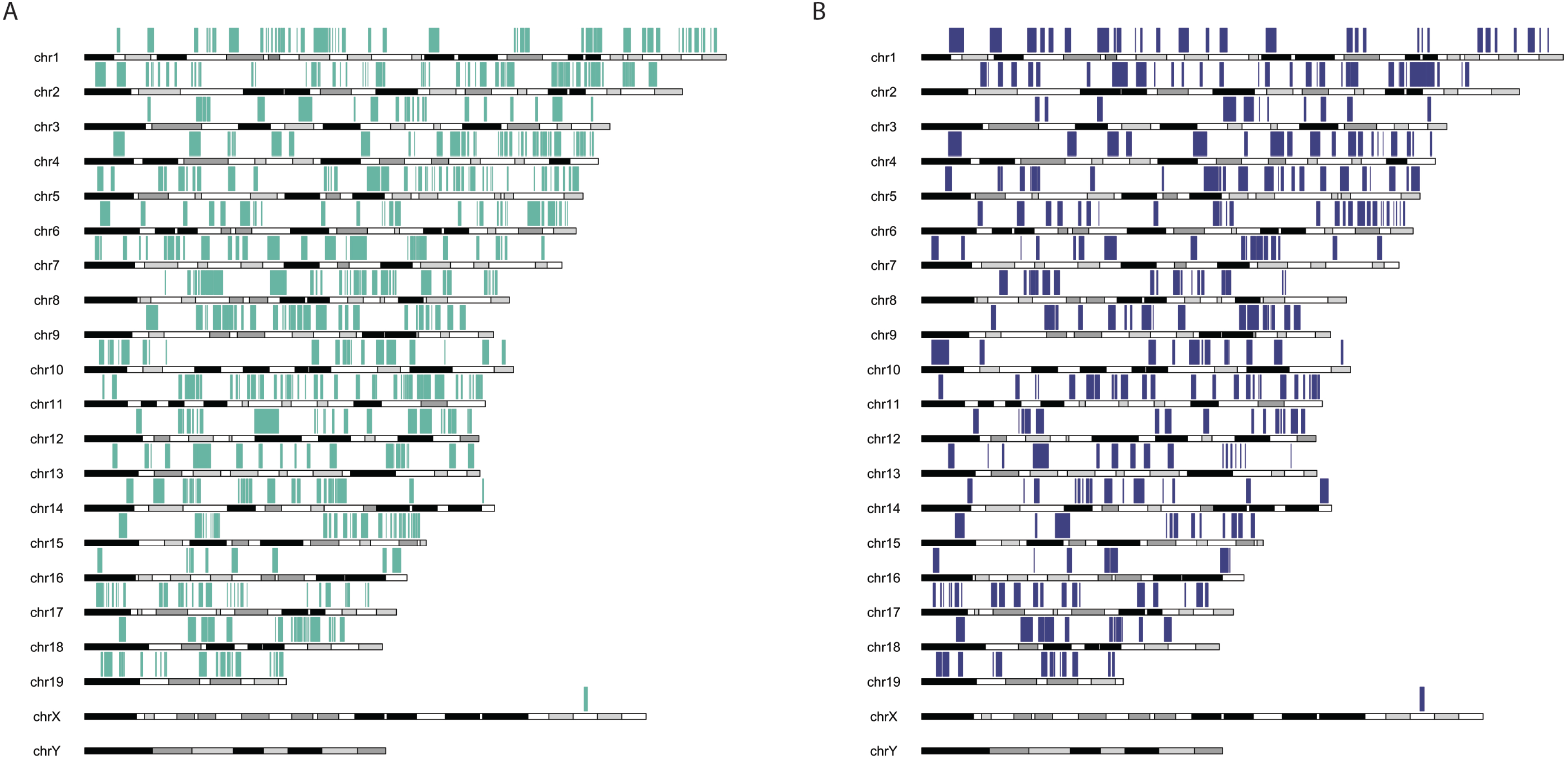
Consensus segments of homozygosity for J:ARC (green) and J:DO (blue). The number of consensus segments of homozygosity in J:ARC samples was higher **(A)** than in J:DO samples **(B)**. Plink’s default parameters were used. Only runs of homozygosity containing at least 100 SNPs, and of total length ≥ 1000 kilobases, are noted.

### Interindividual variability

An important consideration in selecting an outbred laboratory mouse strain and sample size in experimental design is interindividual genetic variability. Taking advantage of our variant data, we used principal component analysis to assess interindividual and interstrain variation in the J:ARC and J:DO. Using all jointly called variants, we found the expected interstrain variation (PC1) but more interindividual variability in J:DO samples than J:ARC (PC2) (Figure 5A). These data show that greater genetic diversity can be achieved with fewer samples in the J:DO by at least an order of magnitude. Unbiased clustering of the J:ARC variant data recapitulated the known pedigree and sibling relationships of the J:ARC samples (Figure 5B), highlighting the utility of variant data for pedigree reconstruction in outbred populations.

**Figure 5.**
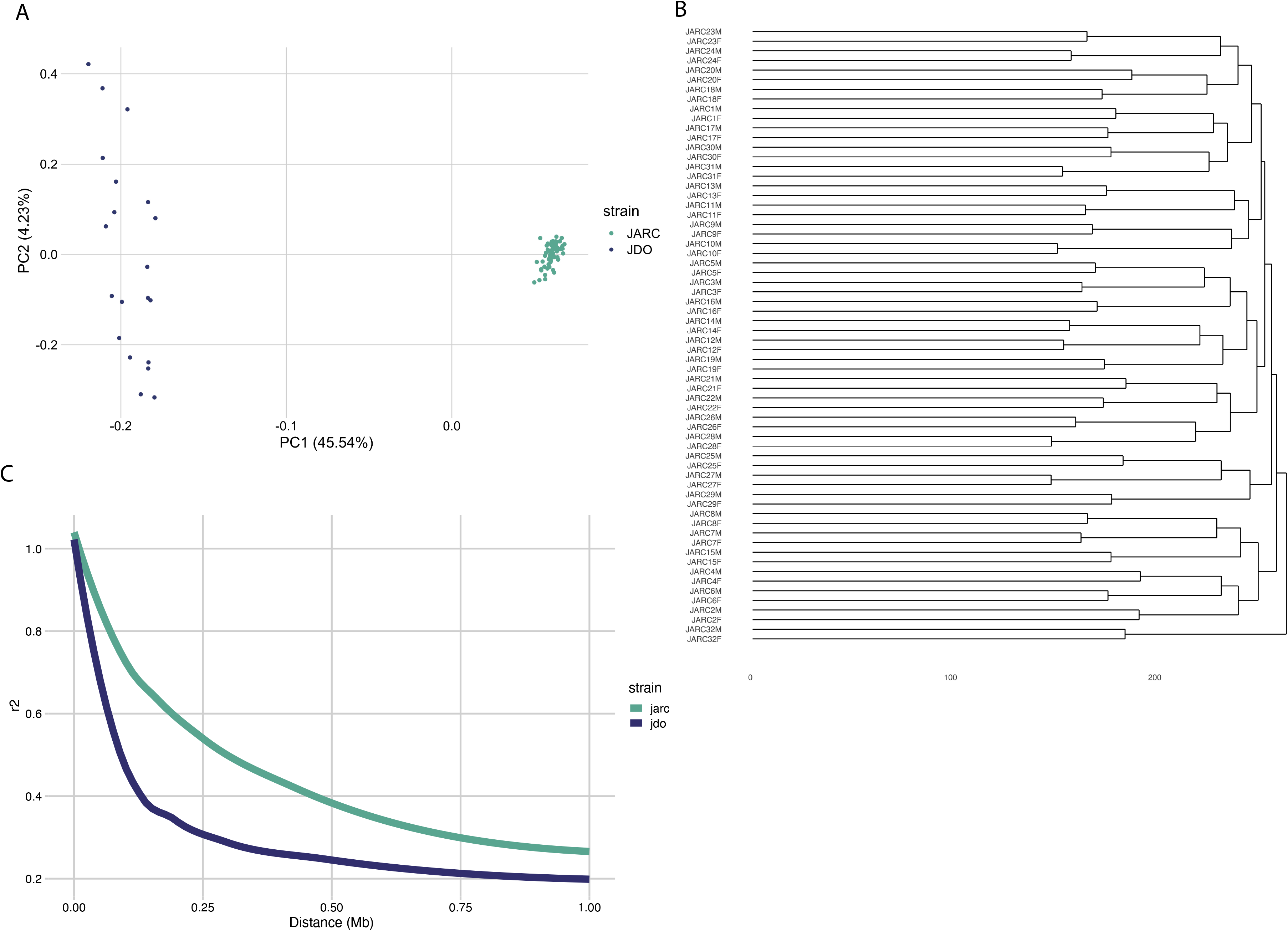
Interindividual variability, relatedness, and LD decay. **(A)** PCA plot of J:DO (blue) and J:ARC (green) joint variant calls. **(B)** A dendrogram of the J:ARC samples (using all called variants in the dataset) showing the samples clustering in known sibling relationships. **(C)** Comparison of LD decay J:ARC samples (green) and J:DO samples (blue). Each curve was plotted using the 95th percentile of r2 values for SNPs spaced up to 1 Mb apart. Plink was used to calculate *r^2^* for all pairs of autosomal SNPs called from joint genotyping in the J:ARC samples (425,409) and the J:DO samples (117,429). SNPs that were missing in more than 5% of the samples and that were monomorphic were removed.

### LD Decay

Linkage disequilibrium (LD) describes the degree to which variants co-segregate in a population. Meiotic recombination leads to new haplotypes and is the molecular mechanism that drives LD decay. Pairwise analysis of linked markers in an interval is used to assess LD decay in a population and these data are useful for inferring swept radius for marker spacing in genetic association studies. Higher LD decay in a mapping population provides higher resolution for genetic mapping. We examined this facet of genetic architecture in the J:ARC and J:DO populations and while our sample size was low, we found that both populations offer LD decay sufficient for genetic mapping, with higher LD decay in the J:DO (LD_1/2_ = ~0.1 Mb) (Figure 5C). Our estimate of LD decay is higher than previous estimates for J:DO, but this is consistent with the more advanced breeding generation used in our study (Gatti *et al*. 2014). LD decay in the J:ARC is consistent with previous estimates for commonly used laboratory outbred populations (LD_1/2_ = ~0.3 Mb) (*Yalcin et al. 2010*).

### Novel Variants

To identify the novel variant calls in the J:ARC and J:DO, we compared the calls to all published variants from all sequenced laboratory mouse strains in dbSNP, EVA, the Sanger mouse genomes project (Lilue *et al*. 2018), Mouse Collaborative Cross Genome (Srivastava *et al*. 2017), and to in house J:DO genome variation call sets (unpublished). Of the 135,825 and 478,011 variants found in J:ARC and J:DO respectively, 96.24 and 99.08% are variants that have been identified in other laboratory mouse strains. We identified 5,105 [2,191 > AF 0.2] and 4,308 [1,888 > AF 0.2] novel variants respectively (Table 1). The larger number of novel variants in J:ARC mice is attributable to the relative lack of published variant data from Swiss derived outbred stocks compared to the complete catalog of variation available from the eight founder inbred strains of the Collaborative Cross recombinant inbred panel from which the J:DO were derived. Functional annotation of the novel variants revealed that J:DO have nearly twice the number of protein-damaging variants (including exon deletions, frameshift, stop gain / loss, splice acceptor / donor sites, start loss) ((J:DO:797 [329 > AF 0.2, HIGH], J:ARC: 408 [151 > AF 0.2, HIGH]) (Figure 6). The details of each variant are provided in Supplementary Data 2. The genes harboring these high impact variants in J:DO show functional enrichment for post-translational modifications (Ubl conjugation, isopepeptide bond), DNA repair, and translational regulation / RNA binding; while the genes harboring high impact variants in J:ARC do not show functional enrichment. The novel protein damaging variants in J:DO are variants that have naturally arisen in the population but have not been lost through purifying selection or through artificial selection. These genetic differences are attributable not only to the distinct origins and ages of these outbred stocks, but to the different breeding paradigms and selective pressures that have occurred during their maintenance. For example, many outbred mouse stocks and populations have been subject to artificial selection for desirable laboratory traits (fecundity, docility, size, etc.) leading to lack of variability for some phenotypes. One example is body weight where variation is less desirable because it complicates pharmacological applications, i.e. dosing studies. As a result, body weight variation in CD-1 and related outbred like J:ARC is minimized, however this is not the case in J:DO where a larger range of body weights is observed (Supplementary Figure 4).

**Figure 6.**
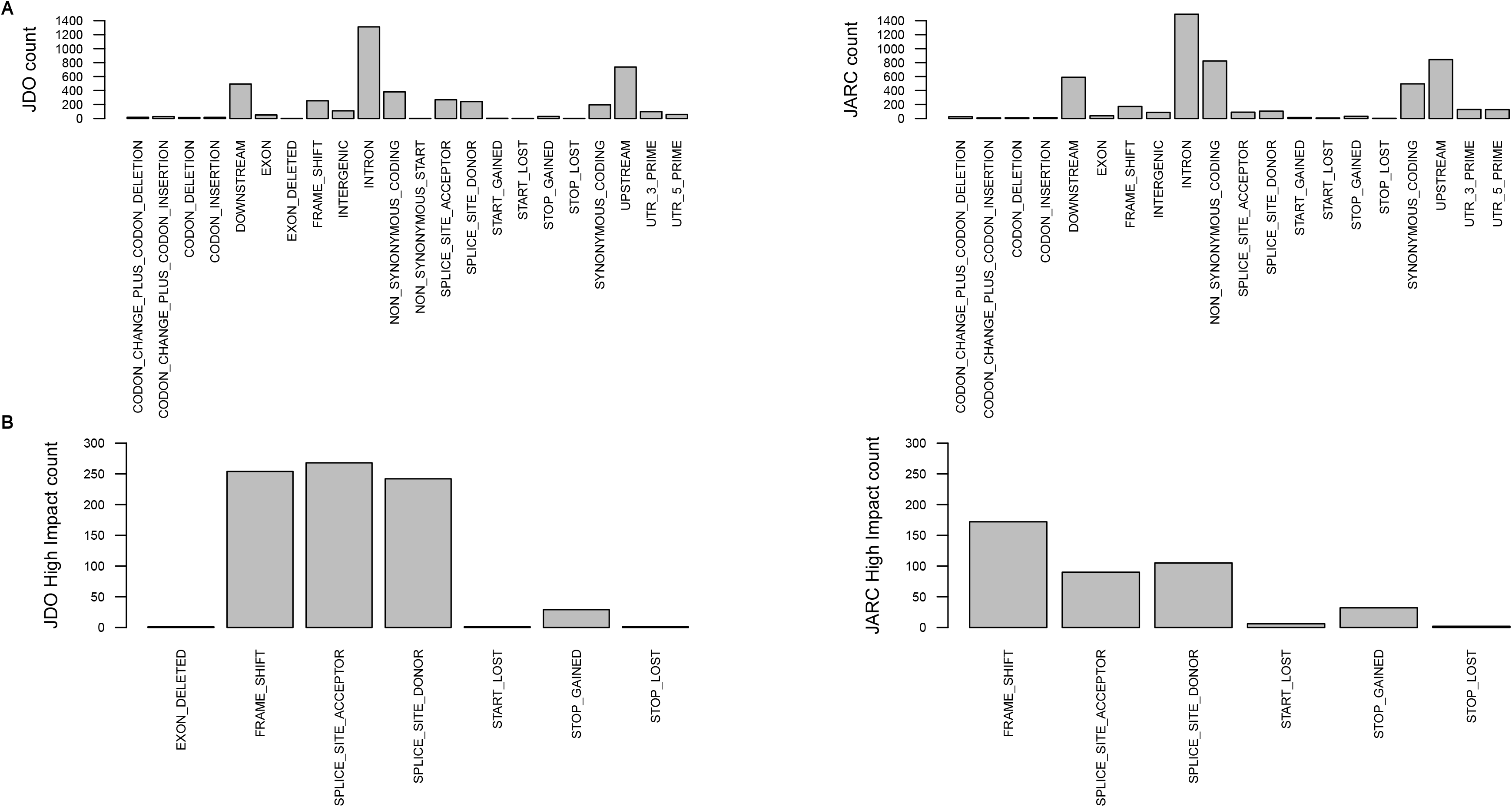
Novel variant consequences. **(A)** Showing the SnpEff effect categories for novel variants for J:DO (on the left) and J:ARC (on the right) **(B)** High-impact (SnpEff effect categories) novel variants are enriched in J:DO (on the left) in comparison to J:ARC (on the right)

**Table 1.** Total variants (SNPs/INDELs) and total novel variants in J:DO and J:ARC stocks. Comparison of the total variants to all variants in the major, publicly available mouse variant resource revealed that 0.9% and 3.75% of the total variants found in J:DO and J:ARC respectively are novel.

| Stock | Total Variants | Mouse Collaborative Cross | Diversity Outbred Mouse Genomes | SangerSNP_Indel_May_2020 | dbSNP | EVA | Novel | High Impact Novel Variants |
| --- | --- | --- | --- | --- | --- | --- | --- | --- |
| J:DO | 478011 | 468487 | 427929 | 452989 | 439682 | 439642 | 4308 | 797 |
| J:ARC | 135825 | 116849 | 109808 | 121395 | 115557 | 115561 | 5105 | 408 |

## CONCLUSIONS

We used exome sequencing to define protein coding variation in two outbred populations of mice with distinct origins. We found that the multiparent, J:DO outbred population has more than 3X population-level variation than the CD-1 derived J:ARC outbred population. Both populations have coefficients of inbreeding that are consistent with previous estimates, confirming that both are maximally outbred. Our analysis of variants that are shared between the J:ARC and J:DO shows that individual heterozygosity in both populations is high, but allelic diversity, intraindividual variability, and minor allele frequency are higher in the J:DO population. For research applications this means that similarly sized cohorts of J:DO mice will provide more genetic diversity than J:ARC, making the former population the better choice for studies that endeavor to maximize genetic diversity. Our data and multiple previous studies show that both outbred populations are useful for genetic mapping. But the higher genome-wide LD decay in J:DO will confer higher mapping resolution.

Overall, we found more novel variants in J:ARC compared to J:DO and this is likely due to the relative paucity of published / accessible variant data for commonly used outbred strains, especially CD-1 from which the J:ARC population was derived. While there were fewer novel variants in the J:DO, more of these were variants that are predicted to be protein damaging and potentially more likely to reduce survival and/or fecundity. While this study doesn’t allow us to make any inferences about the load of deleterious alleles and their respective contributions to fecundity, litter size is a trait that is frequently used to estimate fecundity and breeding data from The Jackson Laboratory show that J:DO females have litter sizes that are comparable to many common laboratory inbred strains (mean litter size = 9.9, https://www.jax.org/strain/009376). The J:ARC population is derived from an outbred stock that has been subject to nearly a century of artificial selection for high fecundity, docility, and other desirable laboratory traits including consistent body weight which is preferred for dosing studies. In contrast, the J:DO are derived from phenotypically diverse recombinant inbred lines (Collaborative Cross) and artificial selection for any phenotype is actively avoided through randomized breeding that optimized for genetic diversity. These differences in selective pressure may also explain the apparent loss of deleterious alleles through drift in the J:ARC. That deleterious alleles have not been subject to purifying selection in the J:DO, also highlights the potential buffering effects of high allelic diversity and heterozygosity.

Our data, together with previously published studies, reinforce the fact that not all outbred stocks have comparable genetic and phenotypic diversity. Direct sequencing of outbred stocks provides molecular basis for genetic quality control, breeding paradigms, and optimization of experimental design for effective deployment these strains in biomedical research.

### Study limitations

Our exome sequencing data provide just a subset of the overall genetic variation segregating in these outbred populations, specifically SNPs and small insertions / deletions in coding sequence. Structural variation (SV), which we have not profiled here, is also a significant source of genetic variation. SV have the potential to impact multiple genes with potentially large phenotypic effects. While certain types of structural variants (deletions) can be robustly detected using short read sequencing data, long read sequencing and *de novo* assembly are the gold standard for genome-wide detection of SV. Moreover, outside of splice junction sites and UTRs, our data do not capture non-coding variation, yet non-coding variation is also a recognized source of phenotypic variation, especially for complex traits. Finally, while our sequencing depth per sample is high, our samples sizes are low, making our study underpowered for detection of ultra-rare SNPs/INDELs.

## Supplementary Material

**Supplementary Data 1.** J:DO and J:ARC samples used for the study.

**Supplementary Data 2.** All novel, protein coding variants in J:ARC and J:DO with functional annotation and allele frequency. Functional annotation (DAVID Bioinformatics Resources) clustering of genes harboring novel variants in J:ARC and J:DO.

**Figure S1.**
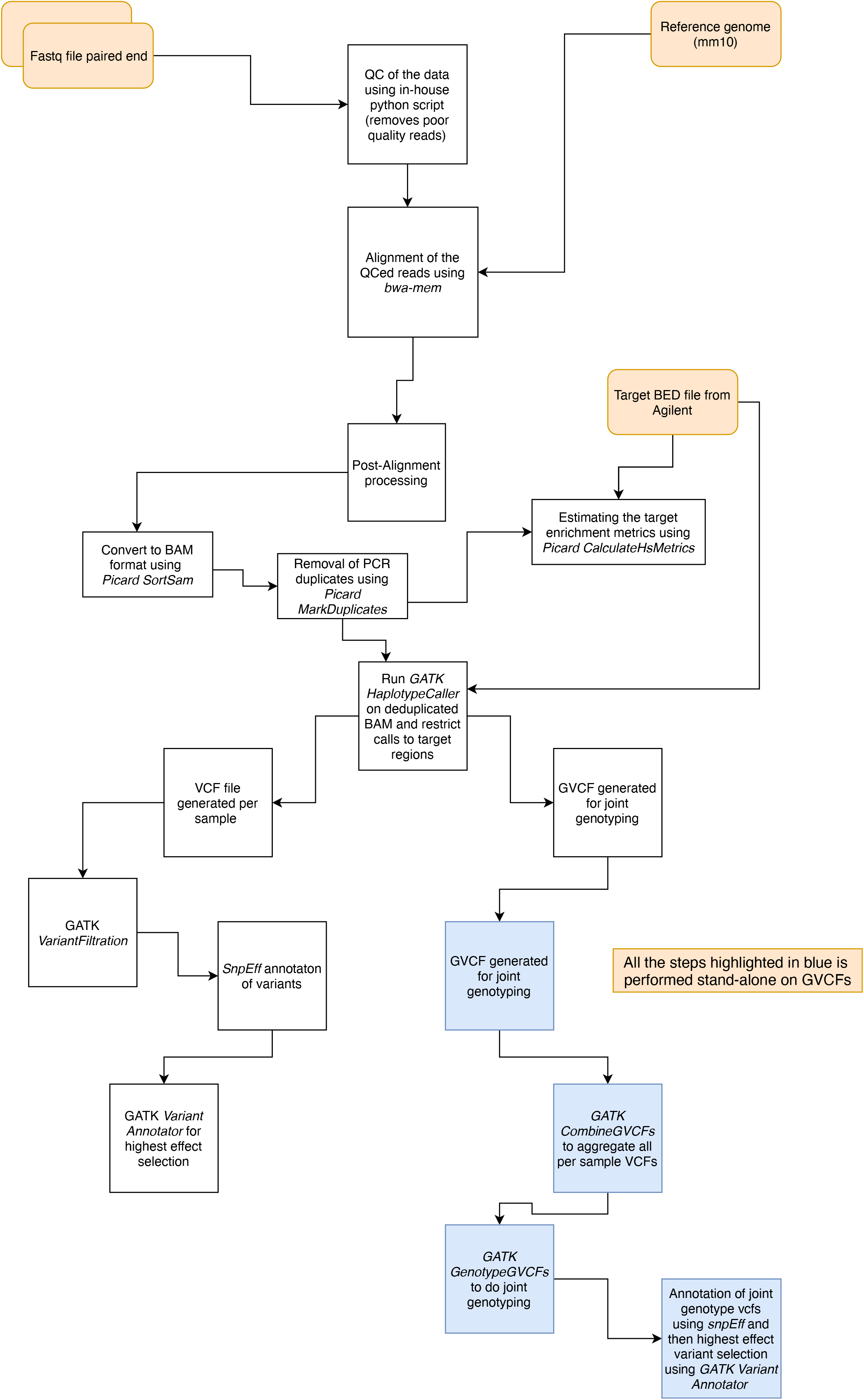
The Jackson Laboratory (JAX) Nexflow pipeline for laboratory mouse whole exome sequencing.

**Figure S2.**
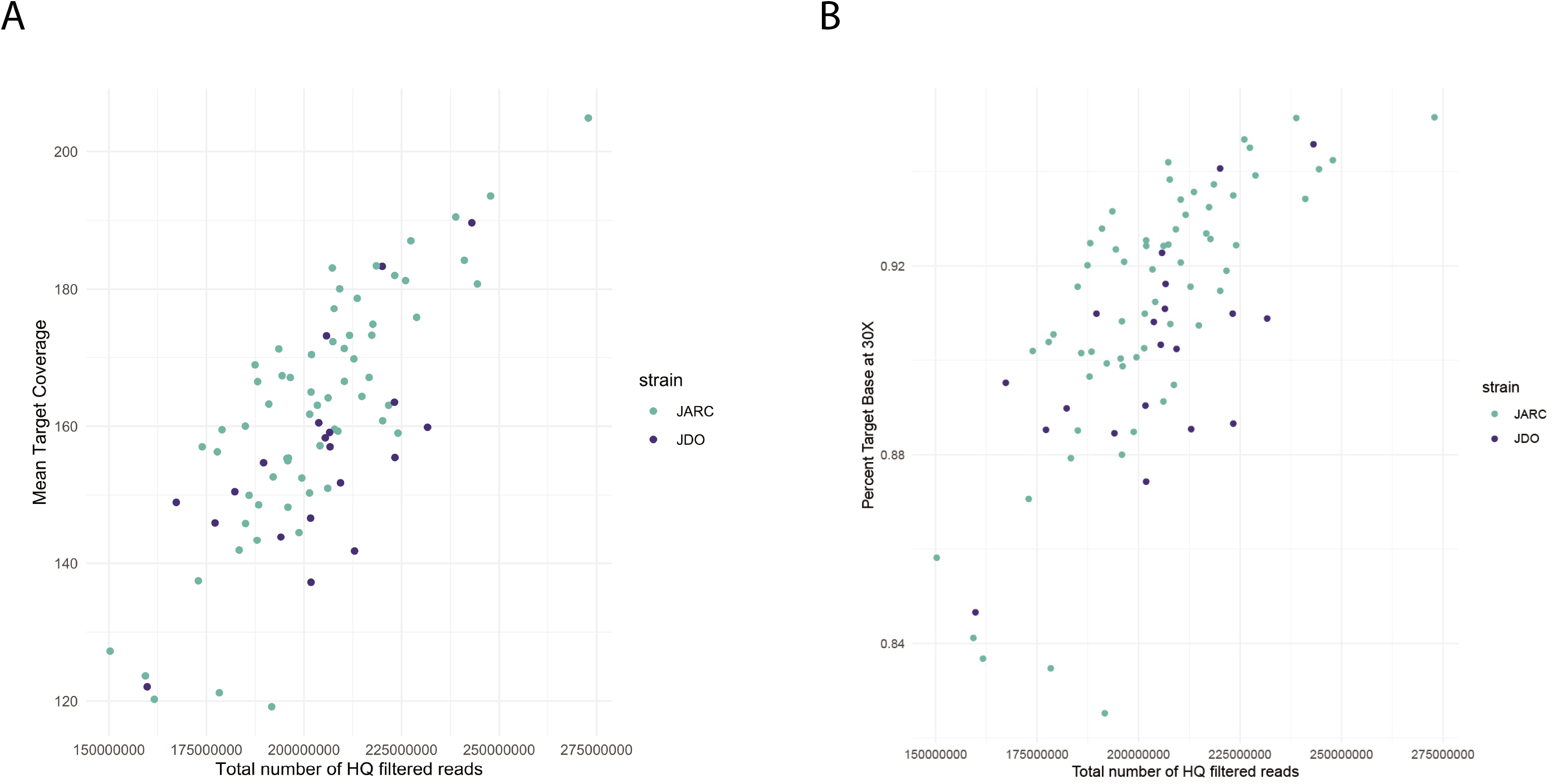
Exome sequencing coverage statistics. **(A)** The mean target coverage versus high quality filtered reads showing mean target coverage was 162X (91% target covered at 30X) for J:ARC samples (green) and 155X (90% target covered at 30X) for J:DO (blue) samples. **(B)** Percent target bases at 30X versus high quality filtered reads showing 90% of the target exome regions were covered by 30 or more reads in both sets of samples.

**Figure S3.**
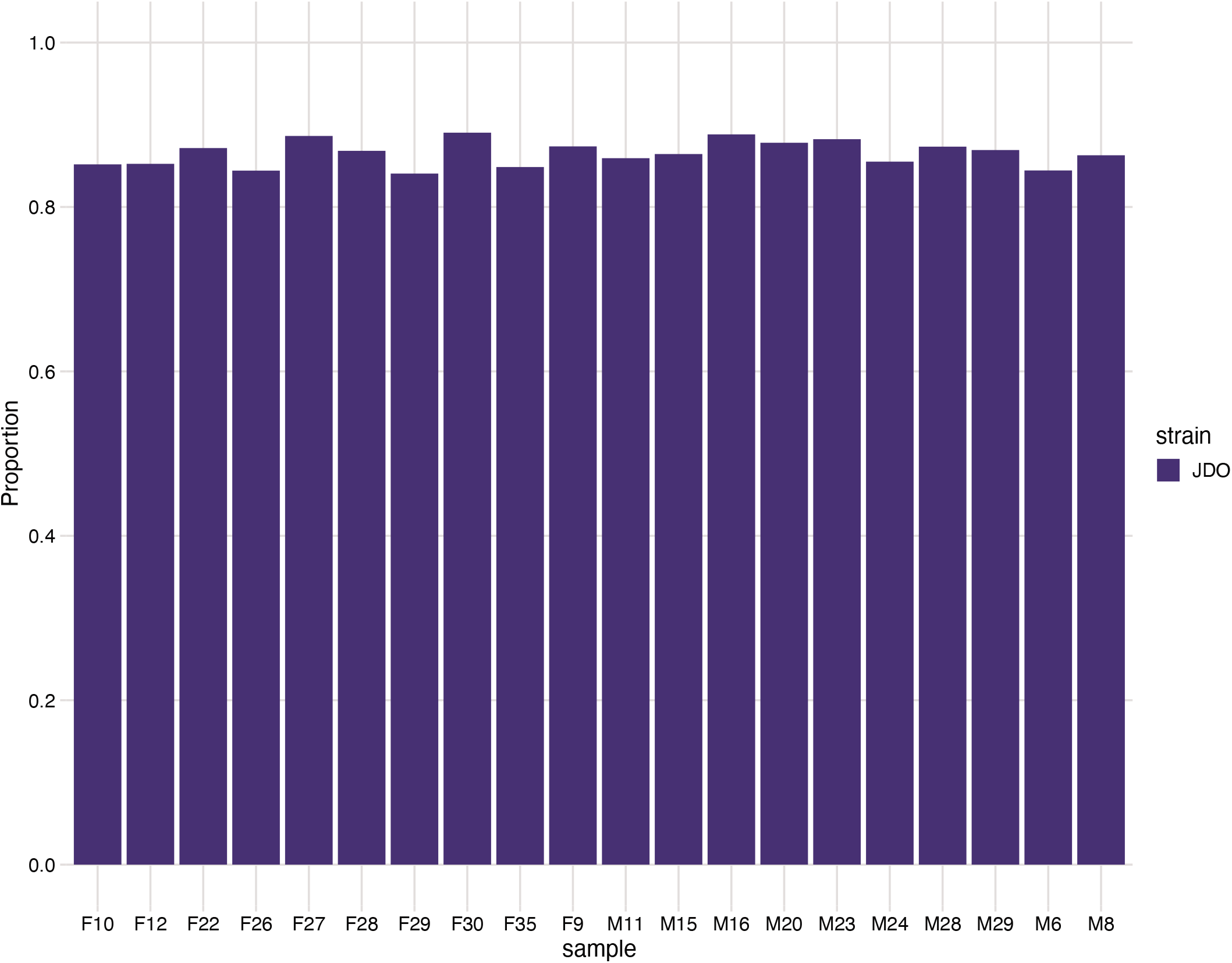
The proportion of heterozygous variants based on the subset of variant calls for known SNPs within the Mouse Universal Genotyping Array probe set. There were 3007 exome variant calls at positions that are included in the GigaMUGA probe set. These were used to estimate the proportion of heterozygous calls per J:DO using the R package, R/qtl2. Estimates are based on autosomes at the haplotype level, using genotype probabilities of the 8 J:DO parental inbred strain genotypes.

**Figure S4.**
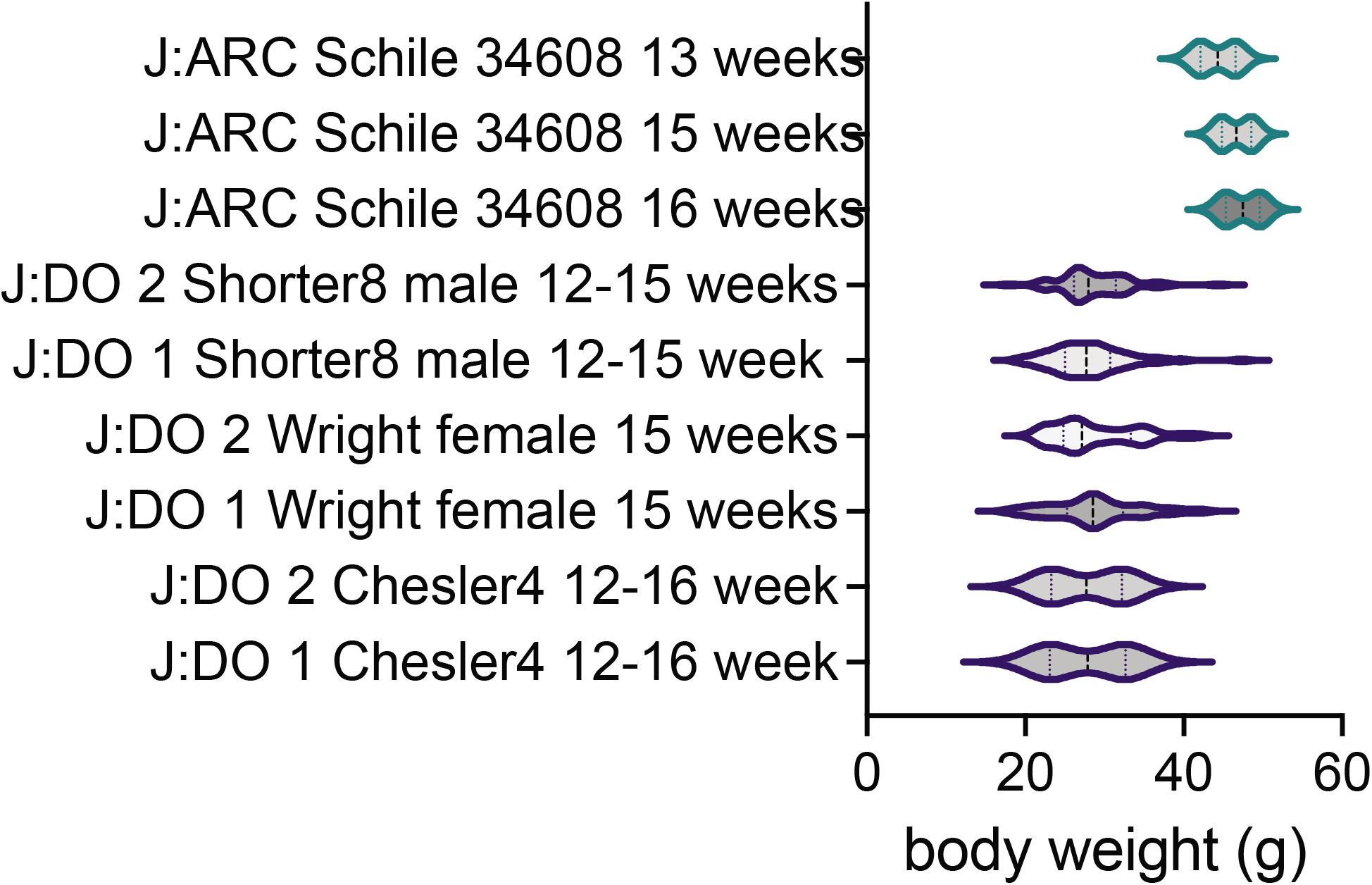
Distribution of body weight in J:ARC and J:DO mice. Jackson Laboratory data (unpublished, Schile 34608) and publicly available datasets from the Mouse Phenome Database ((Shorter8, Chesler4 (Logan *et al*. 2013; Recla *et al*. 2014), https://phenome.jax.org/) and Wright et al. (Wright *et al*. 2022) were used to assess the distribution of body weight in J:DO and J:ARC mice). Data include both males and females and sub-sets of 60 were randomly sampled from larger datasets for comparison. Overall body weight in the J:ARC is higher than J:DO (ANOVA, adjusted P value <0.0001), while the distribution of body weight (range) is greater in J:DO.

## Acknowledgements

This work was partially supported by The Jackson Laboratory and the Mouse Mutant Resource and Research Center at The Jackson Laboratory (NIH U42 OD010921, LGR). We gratefully acknowledge the contribution of the Genome Technologies Service at The Jackson Laboratory for expert assistance with exome library preparation and sequencing.

## Author Contributions

Conceptualization K.B, L.R., A.Sr.

Methodology A.S., B.C.

Formal analysis A.S., B.C.

Investigation A.S., B.C.

Resources C.P., E.S., A. Sh.

Data Curation A.S., B.C.

Writing, Review & Editing L.R., A.Sr., B.C., K.B.

Visualization B.C., A.Sr., L.R.

Supervision A.Sr., L.R.

Funding acquisition L.R., K.B., A.Sr.

## Data availability

All the sequence data used in this study has been submitted to NCBI Sequence Read Archive under the bioproject PRJNA835415 and study SUB11284953.

